# Type 2 Diabetes Modifies Skeletal Muscle Gene Expression Response to Gastric Bypass Surgery

**DOI:** 10.1101/2021.01.19.427352

**Authors:** Matthew D. Barberio, G. Lynis Dohm, Walter J. Pories, Natalie A. Gadaleta, Joseph A. Houmard, Evan P. Nadler, Monica J. Hubal

**Affiliations:** Center for Genetic Medicine Research, Children’s National Medical Center, Washington, D.C.; Department of Exercise and Nutrition Sciences, Milken Institute School of Public Health, George Washington University, Washington, D.C.; Department of Physiology, Brody School of Medicine, East Carolina University, Greenville, N.C.; Department of Surgery, Brody School of Medicine, East Carolina University, Greenville, N.C.; Human Performance Laboratory, Department of Kinesiology, College of Health and Human Performance, East Carolina University, Greenville, N.C.; Division of Pediatric Surgery, Children’s National Medical Center, Washington, D.C.; Department of Kinesiology, Indiana University Purdue University Indianapolis, Indianapolis, IN

## Abstract

Roux-en-Y gastric bypass (RYGB) is an effective treatment for type 2 diabetes mellitus (T2DM) which can result in remission of clinical symptoms, yet mechanisms for improved skeletal muscle health are poorly understood. We sought to define the impact of existing T2DM on RYGB-induced muscle transcriptome changes.

**Methods:** Vastus lateralis biopsy transcriptomes were generated pre- and 1-yr post-RYGB in black adult females with (T2D; n = 5, age=51±6 yr, BMI=53.0±5.8 kg/m^2^) and without (CON; n = 7,43±6 yr,51.0±9.2 kg/m^2^) T2DM. Insulin, glucose, and HOMA-IR were measured in blood at the same time points. ANCOVA detected differentially expressed genes (p< 0.01, Fold change<|1.2|), which were used to identify enriched biological pathways.

**Results:** Pre-RYGB, 95 probes were downregulated with T2D including subunits of mitochondrial complex I. Post-RYGB, the T2D group had normalized gene expression when compared to their non-diabetic counterparts with only 3 probes remaining significantly different. In the T2D, we identified 52 probes upregulated from pre- to post-RYGB, including NDFUB7 and NDFUA1.

**Conclusion:** Black females with T2DM show extensive down regulation of genes across aerobic metabolism pathways prior to RYGB, which resolves 1 year post-RYGB and is related to improvements in clinical markers. These data support efficacy of RYGB for improving skeletal muscle health, especially in patients with T2DM.

## Introduction

Approximately 9 in 10 individuals with type 2 diabetes mellitus (T2DM) are classified as overweight or obese and display peripheral insulin resistance (1). Roux-en Y gastric bypass (RYGB) weight-loss surgery is recognized as an effective intervention for the treatment and remission of T2DM in individuals with severe obesity (2; 3). As the primary site of glucose disposal in response to acute insulin action (4), skeletal muscle is likely a site of improved metabolic programming in response to intervention such as RYGB surgery (5).

Reduced skeletal muscle mitochondrial content and function in response to nutrient oversupply is a significant modifier of skeletal muscle insulin sensitivity in obesity and T2DM (6). Excess intracellular lipids and dysfunctional insulin receptor signaling leads to blunted expression of mitochondrial genes, including the master mitochondrial biogenesis transcriptional regulator peroxisome proliferator-activated receptor co-activator 1 alpha (PCG1α) (7–10). Improved mitochondrial function has been noted in humans following RYGB (11–13), but a limited number of studies have explored skeletal muscle gene expression profiles as far as one year post surgery (5; 14; 15).

Previous studies exploring human skeletal muscle gene expression profiles following RYGB have identified changes in the expression of genes involved in insulin signaling (6), and inflammation (14) as well as mitochondrial and lipid metabolism (15). However, none of these studies specifically address the presence of T2DM on skeletal muscle gene expression compared to people with obesity but not overt T2DM. Previous studies have largely focused on Caucasian subjects despite Black individuals accounting for ~15% of the bariatric surgery population (16). Further, T2DM is more prevalent in Blacks in comparison to other races, underlining the need to study this population in greater detail to understand potential molecular drivers of this disparity. In the current study, we report the effects of T2DM on vastus lateralis global gene expression profiles prior to and one year following RYGB in black women with and without T2DM to determine the modifying effects of existing T2DM in RYGB-response.

## Methods

### Subjects & Clinical Data Collection

Adult Black females without (CON; n = 7) and with T2DM (T2D; n = 5) were recruited from an established bariatric surgery program at Vidant Medical Center (Greenville, NC); all subjects were classified with obesity. Institutional review boards at both East Carolina University and Children’s National Medical Center approved the study and written informed consent was obtained from all study participants. Criteria for inclusion included: age between 25 and 60 years, BMI between 35 and 65 kg/m^2^, a negative pregnancy test, and (for T2D group) a diagnosis of T2DM in accordance with the criteria for the NIH Consortium for the Longitudinal Assessment of Bariatric Surgery (17).

Subjects were in enrolled in the standard clinical protocol for bariatric surgery at Vidant Health. Anthropometric measures (age, height, weight, BMI), fasting blood (antecubital) collection, and skeletal muscle biopsies were collected two weeks prior to (Pre) and 1-year post-surgery (Post). Whole blood was collected in plasma and serum separating tubes. BMI was calculated as kg/m^2^ and percent excess BMI loss (% Excess BMI Loss) calculated as ((Pre BMI – Post BMI) / (Pre-BMI – 25)) x 100. Insulin was measured by immunoassay (Access Immunoassay System, Beckman Coulter, Fullerton, CA) and glucose with an oxidation reaction (YSI 2300, Yellow Springs, OH). The homeostasis model assessment (HOMA2) was calculated from plasma glucose and insulin levels (www.dtu.ox.ac.uk/homacalculator) (18).

### Skeletal Muscle Biopsies

Skeletal muscle biopsies were taken two weeks prior to and 1-year following RYGB, in a fasted state, from the vastus lateralis muscle of the non-dominant leg via standard Bergstrom needle biopsy(19). Approximately 100 to 200 mg of tissue was obtained with a triple pass and immediately flash frozen in liquid nitrogen. Approximately 20 to 30 mg of tissue was used for both RNA and DNA extractions protocols

### Gene Expression Profiling

Skeletal muscle gene expression was analyzed from skeletal muscle biopsies taken Pre- and Post-RYGB via global microarray analysis (Affymetrix HU133 Plus 2.0 microarray; Affymetrix, Santa Clara, CA; Accession: GSE161643). Total RNA was isolated from skeletal muscle homogenates via the TRIzol (Invitrogen. Carlsbad, CA) method (20). Affymetrix instructions were followed for microarray processing. Briefly, 500ng of total RNA was used with appropriate Poly-A controls for first- and second-strand cDNA synthesis. Biotin labeled complementary RNA (cRNA) was synthesized using *in vitro* transcription of the second-strand cDNA with a T7 RNA polymerase. Approximately 30 ug of labeled cRNA was fragmented and hybridized to each microarray.

CEL files were generated from scanned microarrays and imported into Affymetrix Expression Console, where CHP files were generated using the PLIER (Probe Logarithmic Intensity Error) algorithm. PLIER is a mode-based signal estimator which takes advantage of numerous internal control probes of the microarray to differentiate between background and signal. Standard quality control methods were used to evaluate amplifications, thresholds for appropriate scaling factors, and RNA integrity (GAPDH 3’/5’ and HSAC07 3’/5’). Samples failing quality standards were reprocessed from original total RNA. Probe lists for statistical analysis (using the PLIER generated probe intensities) were also filtered for present/absent calls using the MAS5.0 algorithm in Expression Console. Probes that were determined present on 20 of 24 arrays (83.3%) were retained for statistical analysis. Resultant CHP files were imported into Partek Genomics Suite (Partek, Inc.; St. Louis, MO). Probe set intensities (PLIER) were log_2_-transformed for data normalization before statistical analyses.

### Differential Expression Analysis and Biological Interpretation of Gene Expression

Differences in gene expression were assessed via 3-Way Analysis of Covariance using a restricted maximum likelihood approach (Model: timepoint x group x timepoint*group + age + BMI) with contrasts between groups and timepoints conducted via Fisher’s Least Significant Difference test (21). Significant probes were defined as p < 0.01 and resultant gene sets were uploaded to Ingenuity Pathway Analysis (IPA; Qiagen, Inc.) for probe set annotations and to query relationships between genes. The canonical pathway analysis tool was used to identify biological pathways that were overrepresented in our data set via a Right-Handed Tukey’s T-Test (22). We also utilized Gene Ontology (GO) Enrichment Analysis to determine categorization of the biological processes of significant gene lists (23; 24). GO Enrichment Analysis uses a Fisher’s Exact test for classification and p-values for False Discovery Rate (FDR) are reported in the results.

### Real-Time PCR Validation of Target Genes

Microarray results were confirmed with real-time polymerase chain reaction (qPCR). Due to RNA quantity and concentrations available following microarray analysis, a representative subset of n = 3 in the T2D group was used in qPCR analysis. RNA (100 ng) was reverse-transcribed into cDNA using SuperScript III Reverse Transcription (Invitrogen Corp.; Carlsbad, CA) following manufacturer protocols. PCR was performed in triplicate on an Applied Biosystems QuantStudio 3 Real-Time PCR Systems with Taqman Universal PCR Master Mix and commercially available TaqMan human gene expression assays (ThermoFischer Scientific; Waltham, MA) for hexokinase 2 (HK2; AssayID: Hs00606086_m1), NADH:ubiquinone oxidoreductase subunit B8 (NDUFB8; Hs00428204_m1), NADH:ubiquinone oxidoreductase subunit B7 (NDUFB7; Hs00958815_g1), NADH:ubiquinone oxidoreductase subunit A1 (NDUFA1; Hs00244980_m1), 3-hydroxybutyrate dehydrogenase, type 1 (BDH1; Hs00366297_m1). Assays were performed in accordance with manufacturer instructions: 50°C for 2 min, 95° for 10 min, followed by 40 cycles of 95°C for 15 sec followed by 60°C for 1 min. Assays were run with a multiplexed endogenous control (B2M). Fold changes were determined via the 2^-ΔΔCt^ methodology.

### Statistical Analyses

Transcriptome and pathway statistical analyses are described above. Clinical data normality was assessed with Shapiro-Wilk tests and visualization of the distribution. If data were non-normally distributed, the data were log_2_-transformed and reassessed for normality. Differences between groups for anthropometric and clinical data were tested via Two-Sample T-Test (age, height, % excess BMI Loss, % HOMA-2 Change) and Two-Way Repeated Measure ANOVA (Group x Time x Group*Time) for remaining measures. When appropriate, a Tukey’s Test was used for pairwise comparison. Statistical analyses were performed using OriginLab Pro 2015 (OriginLab Corp, Northampton, MA).

## Results

### Clinical Characteristics

Anthropometric and clinical characteristics are presented in Table 1. The T2D group was significantly (p = 0.03) older than subjects without diabetes. A time*group interaction (p = 0.03) was observed for changes in BMI. Analysis of percent of excess BMI loss indicated CON lost significantly (p =0.03) more excess BMI as compared to T2D. Blood glucose was significantly (p = 0.04) reduced in both groups 1-year following surgery with no significant (p = 0.06) difference between groups either pre- or post-surgery. A time x group interaction (p < 0.001) was observed for blood insulin. Post-hoc analysis identified that blood insulin was significantly reduced in T2D (p = 0.03) and CON (p = 0.01) diabetes following surgery. Furthermore, CON had significantly lower blood insulin pre (p = 0.007) and post (p = 0.01) surgery as compared to T2D. Main effects for time (p = 0.001) and group (p < 0.001) were observed for HOMA-IR indicating CON had lower HOMA-IR pre and post-surgery though both groups significantly improved following surgery. However, no difference (p=0.70) was detected between groups for percent change in HOMA-IR following surgery.

**Table 1.**
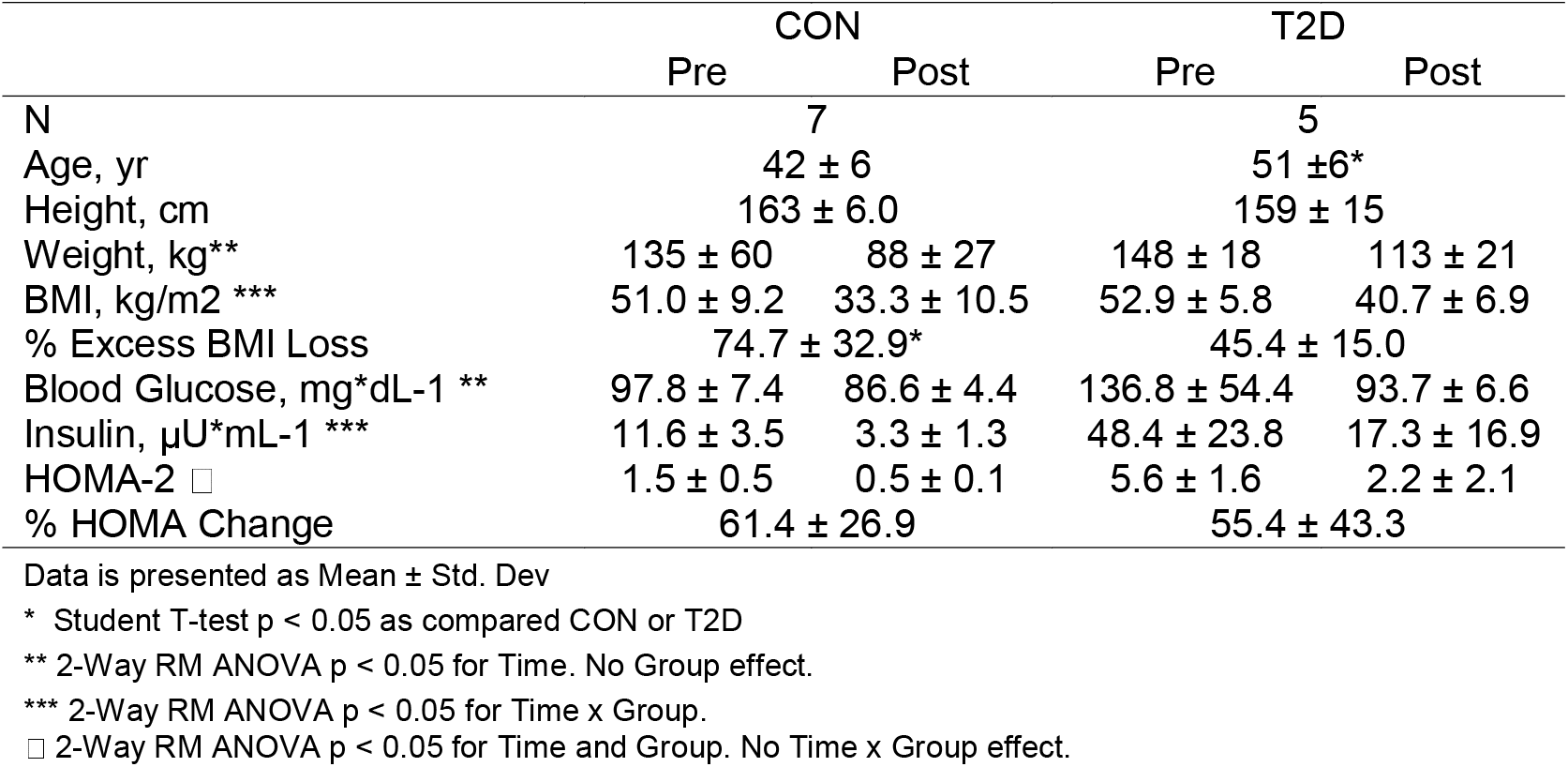
Clinical Characteristics.

|  | CON |  | T2D |  |
| --- | --- | --- | --- | --- |
|  | Pre | Post | Pre | Post |
| N | 7 |  | 5 |  |
| Age, yr | 42 ± 6 |  | 51 ± 6* |  |
| Height, cm | 163 ± 6.0 |  | 159 ± 15 |  |
| Weight, kg** | 135 ± 60 | 88 ± 27 | 148 ± 18 | 113 ± 21 |
| BMI, kg/m2 *** | 51.0 ± 9.2 | 33.3 ± 10.5 | 52.9 ± 5.8 | 40.7 ± 6.9 |
| % Excess BMI Loss | 74.7 ± 32.9* |  | 45.4 ± 15.0 |  |
| Blood Glucose, mg*dL-1 ** | 97.8 ± 7.4 | 86.6 ± 4.4 | 136.8 ± 54.4 | 93.7 ± 6.6 |
| Insulin, µU*mL-1 *** | 11.6 ± 3.5 | 3.3 ± 1.3 | 48.4 ± 23.8 | 17.3 ± 16.9 |
| HOMA-2 □ | 1.5 ± 0.5 | 0.5 ± 0.1 | 5.6 ± 1.6 | 2.2 ± 2.1 |
| % HOMA Change | 61.4 ± 26.9 |  | 55.4 ± 43.3 |  |

Data is presented as Mean ± Std. Dev
\* Student T-test $p < 0.05$ as compared CON or T2D
\*\* 2-Way RM ANOVA $p < 0.05$ for Time. No Group effect.
\*\*\* 2-Way RM ANOVA $p < 0.05$ for Time x Group.
□ 2-Way RM ANOVA $p < 0.05$ for Time and Group. No Time x Group effect.

### Skeletal Muscle Global Gene Expression

Following preliminary filtering for MAS5.0 present/absent (83.3% cutoff) left 25,839 probes for statistical analysis using PLIER generated probe intensity values. ANCOVA detected 114 significant (p < 0.01; **Supplementary Table 1**) probes for the main effect of timepoint, 61 significant (p < 0.01; **Supplementary Table 2**) probes for the main effect of group, and 376 significant (p < 0.01; **Supplementary Table 3**) probes for time x group interaction. The 376 significant probes from the time x group interaction were carried forward for pairwise comparisons.

### Pre-Surgery Differences in Skeletal Muscle Oxidative Metabolism Gene Expression

Baseline pairwise comparison (Pre-RYGB T2DM vs Pre-RYGB CON) resulted in 97 significant probes (**Figure 1A**, **Supplemental Table 4**) of which 95 had lower expression in T2D. We used multiple bioinformatics knowledge base tools to explore the biological context of our significant probe list. Using Gene Ontology Enrichment Analysis to categorize our probes based on biological processes, 82 of the 97 probes mapped to known genes and 52 of those categorized under GO:0008152 Metabolic Process (FDR p = 1.72 x 10^−2^). **Figure 1B** depicts the most significant Gene Ontology terms from significant genes prior to surgery. IPA identified 11 enriched canonical biological pathways (**Figure 1C**), the majority of which are involved in oxidative metabolism. The top canonical pathways identified were “Mitochondrial Dysfunction” (p = 1.66 x 10^−11^; 12 genes), “Oxidative Phosphorylation” (p =4.45 x 10^−11^; 10 genes), and “Sirtuin Signaling Pathway” (8.89 x 10^−8^; 11 genes). Genes involved in oxidative metabolism identified via Canonical Pathway Analysis in IPA are listed **Table 2.** Differences in gene expression between groups pre-surgery for 5 genes (**Figure 1D**) were confirmed via qPCR.

**Table 2.**
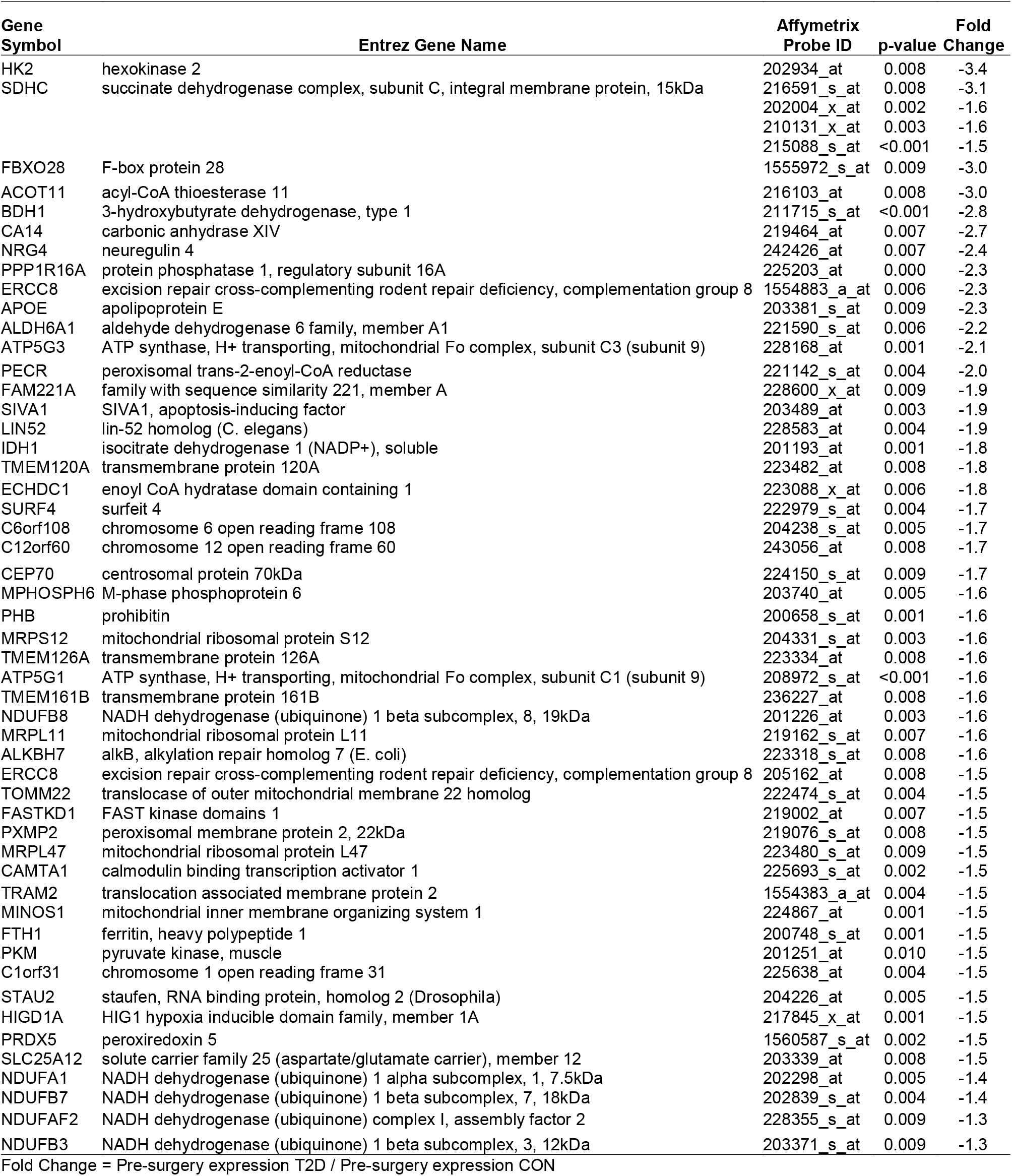
Baseline (Pre-RYGB) group differences in skeletal muscle gene expression for oxidative pathways.

**Figure 1.**
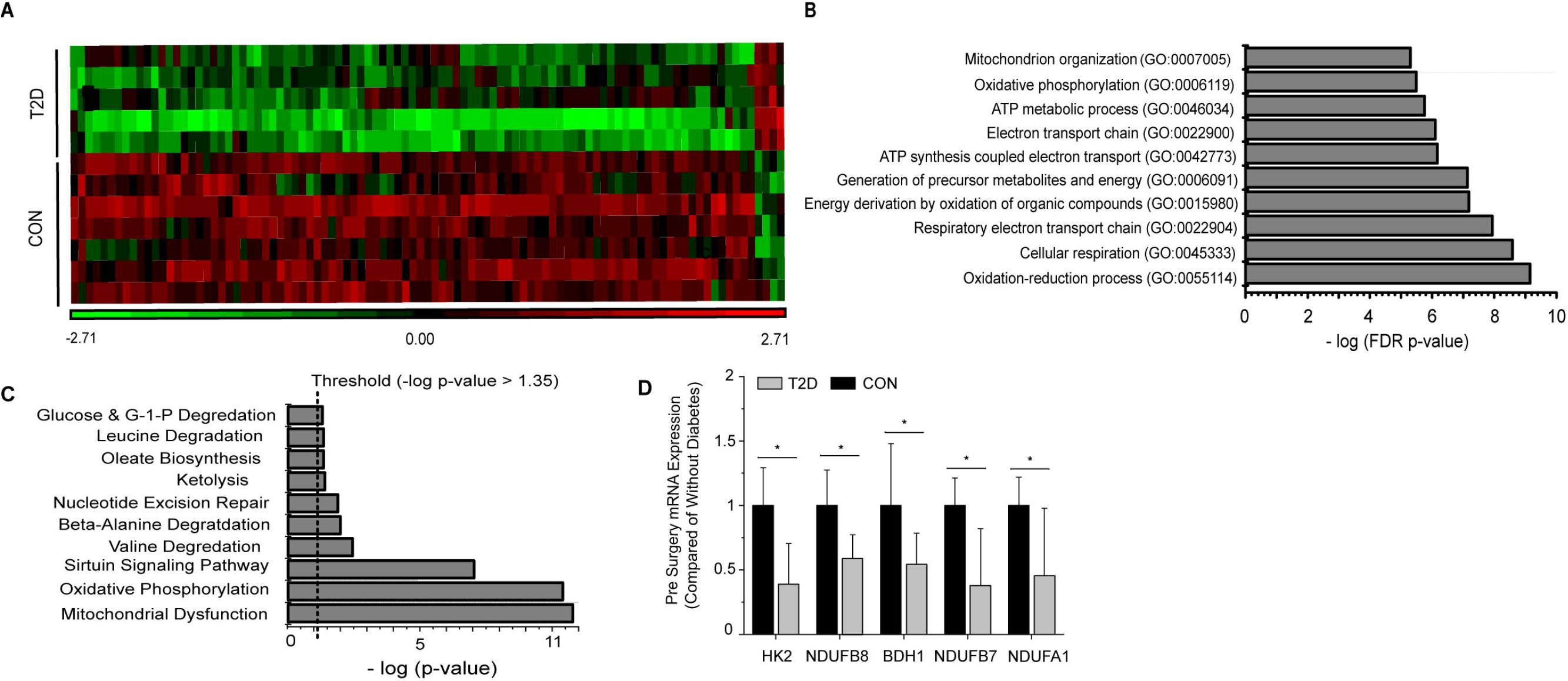
Pre-surgery skeletal muscle gene expression. **A)** Heat map (probe intensities) of 97 significantly (p < 0.01; FC > |1.2|) probes used for biological function and pathway analysis. **B)** Gene Ontology Enrichment analysis for biological process of differentially expressed genes in individuals with and without diabetes pre-surgery. **C)** Significant pathways from Ingenuity Pathway Analysis using the Canonical Pathway tool. Presented pathways are overrepresented with genes differentially regulated between those with and without diabetes. **D**) Fold changes in select genes as determined via qPCR; * p < 0.05 students t-test.

### Improvements in Oxidative Metabolism Gene Expression and Resolution of Gene Expression Differences 1-Year Post Surgery in Individuals with Diabetes

Comparison of skeletal muscle gene expression profiles 1-year post surgery resulted in only 3 significant probes which mapped to 3 genes: ring finger protein 6 (RNF6; p = 0.003, FC = 1.8 greater in individuals with diabetes vs. without), coiled-coil domain containing 90A (CCDC90A; p = 0.007, FC = 1.3), and guanine monophosphate synthetase (GMPS; p=0.002, FC = −1.3). These data represent a “closing of the baseline gap” between T2D and CON skeletal muscle health post-surgery. Only CCDC90A (p = 0.002; FC = −1.2) was found to be significantly different in pre-surgery gene expression analysis.

We explored changes in skeletal muscle gene expression profiles pre-to-post surgery in individuals with diabetes and without diabetes (**Figure 2A**). Comparison of pre-to-post surgery gene expression changes identified 53 probes (**Supplemental Table 5**), mapping to 48 known genes. An increase in expression (FC > 1.2) was observed in 43 of the 48 known genes (**Figure 2B)**. Only 11 genes were significantly altered pre-to-post surgery in individuals without diabetes. Gene Ontology Enrichment Analysis for Biological Process of genes with significant differential expression pre-to-post surgery in individuals with diabetes identified respiratory electron transport chain (FDR = 6.8 x 10^−2^) and oxidative phosphorylation (FDR = 4.18 x 10^−2^) as the only significant biological processes. Similarly, canonical pathway analysis again identified Oxidative Phosphorylation (5.19 x 10^−8^; 6 genes), Mitochondrial Dysfunction (1.15 x 10^−6^; 6 genes), and Sirtuin Signaling Pathway (3.3 x 10^−3^; 4 genes) as the top overrepresented pathways in the gene list (**Figure 2C**). Significant genes pre-to-post surgery involved in oxidative metabolism identified via Canonical Pathway Analysis in IPA are listed **Table 3**. Increased expression of three genes (i.e. fold changes) in individuals with diabetes were confirmed via qPCR, but were not statistically different (**Figure 2D**).

**Table 3.** Surgery-responsive changes in genes in oxidative pathways in T2D group.

| **Gene**<br>**Symbol** | **Entrez Gene Name** | **Affymetrix**<br>**Probe ID** | **p-value** | **Fold Change**<br>**Pre to Post**<br>**Surgery** |
| --- | --- | --- | --- | --- |
| ATP5O | ATP synthase, H<sup>+</sup> transporting, mitochondrial F1 complex, O subunit | 216954_x_at | 0.006 | 1.3 |
| COX6C | cytochrome c oxidase subunit VIc | 201754_at | 0.005 | 1.3 |
| NDUFB7 | NADH dehydrogenase (ubiquinone) 1 beta subcomplex, 7, 18kDa | 202839_s_at | 0.002 | 1.3 |
| UQCR10 | ubiquinol-cytochrome c reductase, complex III subunit X | 218190_s_at | 0.003 | 1.3 |
| PRDX5 | peroxiredoxin 5 | 222994_at | 0.007 | 1.3 |
| NDUFAF2 | NADH dehydrogenase (ubiquinone) complex I, assembly factor 2 | 228355_s_at | 0.007 | 1.3 |
| TUBA4A | tubulin, alpha 4a | 212242_at | 0.003 | 1.4 |
| NDUFA1 | NADH dehydrogenase (ubiquinone) 1 alpha subcomplex, 1, 7.5kDa | 202298_at | 0.006 | 1.4 |
| ATP5G1 | ATP synthase, H<sup>+</sup> transporting, mitochondrial Fo complex, subunit C1 (subunit 9) | 208972_s_at | <0.001 | 1.6 |
| BDH1 | 3-hydroxybutyrate dehydrogenase, type 1 | 211715_s_at | 0.008 | 1.8 |

Fold Change = post-surgery expression T2D / pre-surgery expression T2D

**Figure 2.**
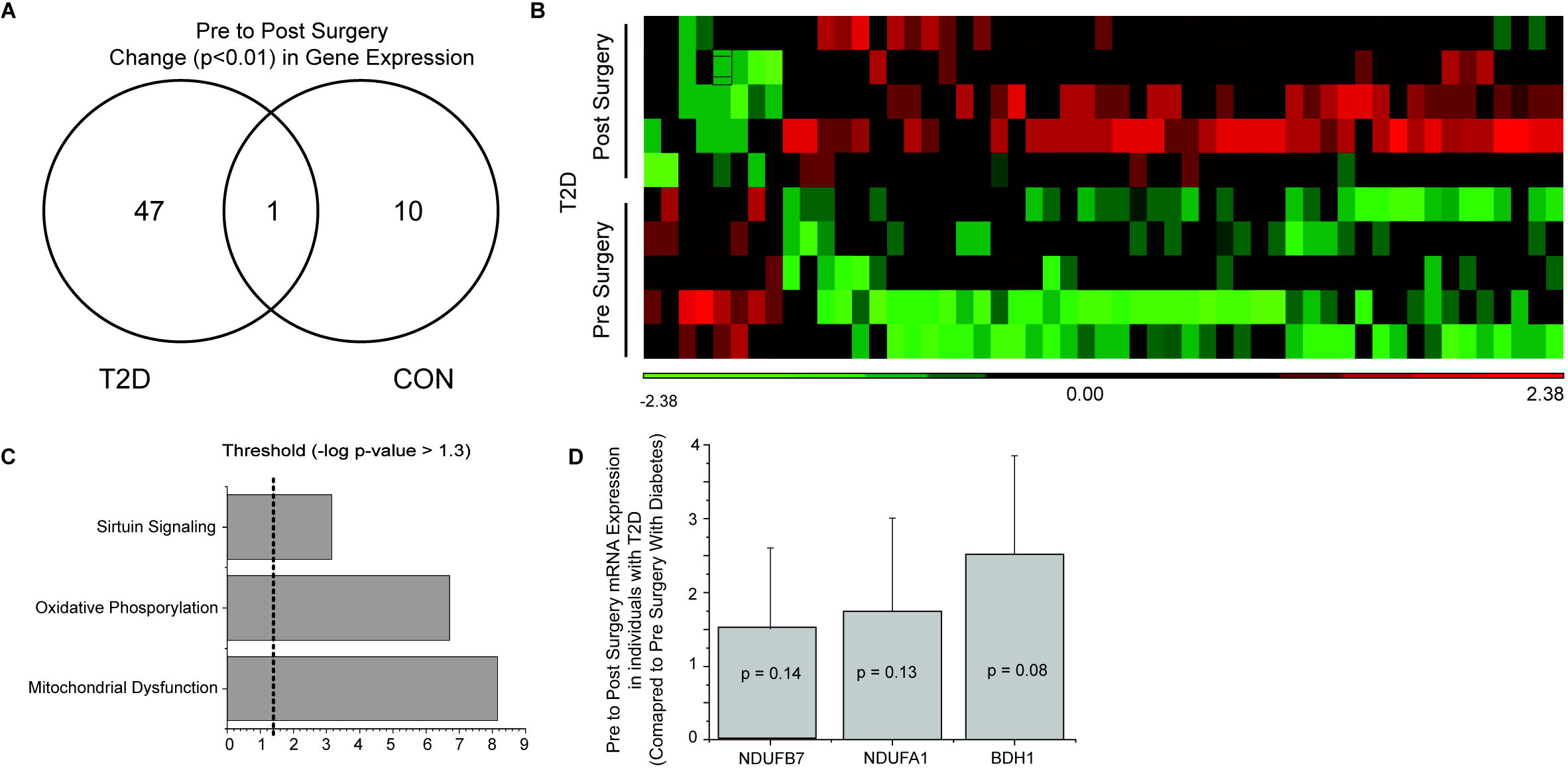
Pre to post-surgery skeletal muscle gene expression. **A)** Venn diagram of significant probes in individuals with diabetes and without diabetes, and overlap, pre-to-post surgery. **B)** Heat map (probe intensities) of 48 significant (p < 0.01, FC > |1.2|) probes pre-to-post surgery in individuals with diabetes. **C)** Significant pathways from Ingenuity Pathway Analysis using the Canonical Pathway tool. Presented pathways are overrepresented with genes differentially regulated between pre-to-post surgery in individuals with diabetes. **D**) Fold changes in select genes as determined via qPCR.

## Discussion

The purpose of the present study was to identify the effect of T2DM on skeletal muscle gene expression profiles before and after RYGB surgery in severely obese Black women. Our data provide evidence that oxidative metabolism genes in women with T2DM are lower prior to RYGB compared to those without T2D, suggesting more dysfunction with the disease. One year following RYGB, skeletal muscle gene expression profiles of those with diabetes normalized compared to their non-diabetic counterparts, largely driven by significant increases in gene expression of electron transport chain genes. Mitochondrial function and perturbations in oxidative metabolism have been implicated in the development of insulin resistance and T2DM (6; 8; 10). RYGB is an effective strategy for significant reductions in excess weight and BMI, as well as the remission of T2DM, though mechanistic understanding of how this resolution occurs remains a work in progress (2; 3). Taken in conjunction with improvements in cardiometabolic profiles (decreased resting blood glucose, insulin, HOMA-2, weight, and BMI), coordinated increases in skeletal muscle oxidative metabolism gene expression appears to play, in part, a role in the remission of T2DM following RYGB surgery.

### Reduced expression of aerobic and mitochondrial pathways in skeletal muscle of Black women with T2D

Individuals with T2DM show significant reductions in skeletal muscle oxidative metabolism gene expression prior to bariatric surgery (15; 25; 26). The current data indicate reductions in multiple subunits of mitochondrial complex I (NDUFB8, NDUFA1, NDUFB7, NDUFAF2, NDUFB3) in individuals with T2DM prior to RYGB (**Figure 3A**). The role of complex I in the development and treatment of T2DM has been explored in various tissue and cells with conflicting conclusions. Chemical inhibition and gene silencing of complex I shows improved glucose consumption in HepG2 and C2C12 cells as well as improved glucose homeostasis in db/db mice (27) whereas complex I deficits result in metabolic inflexibility in the diabetic heart (28). Most recent evidence suggests clinical dosage of metformin improves complex I activity (29) through activation of AMPK despite previous reports of supraphysiologic dosages causing inhibition (30). There is also evidence that complex I deficits restrict fetal skeletal muscle growth (31) and decrease mitochondrial efficacy in aging skeletal muscle (32). Expanded analysis (Supplementary Table 6) of skeletal muscle gene expression differences in those with and without diabetes prior to surgery shows consistent down regulation of a number of genes for mitochondrial complexes in individuals with diabetes (**Figure 3A**).

**Figure 3.**
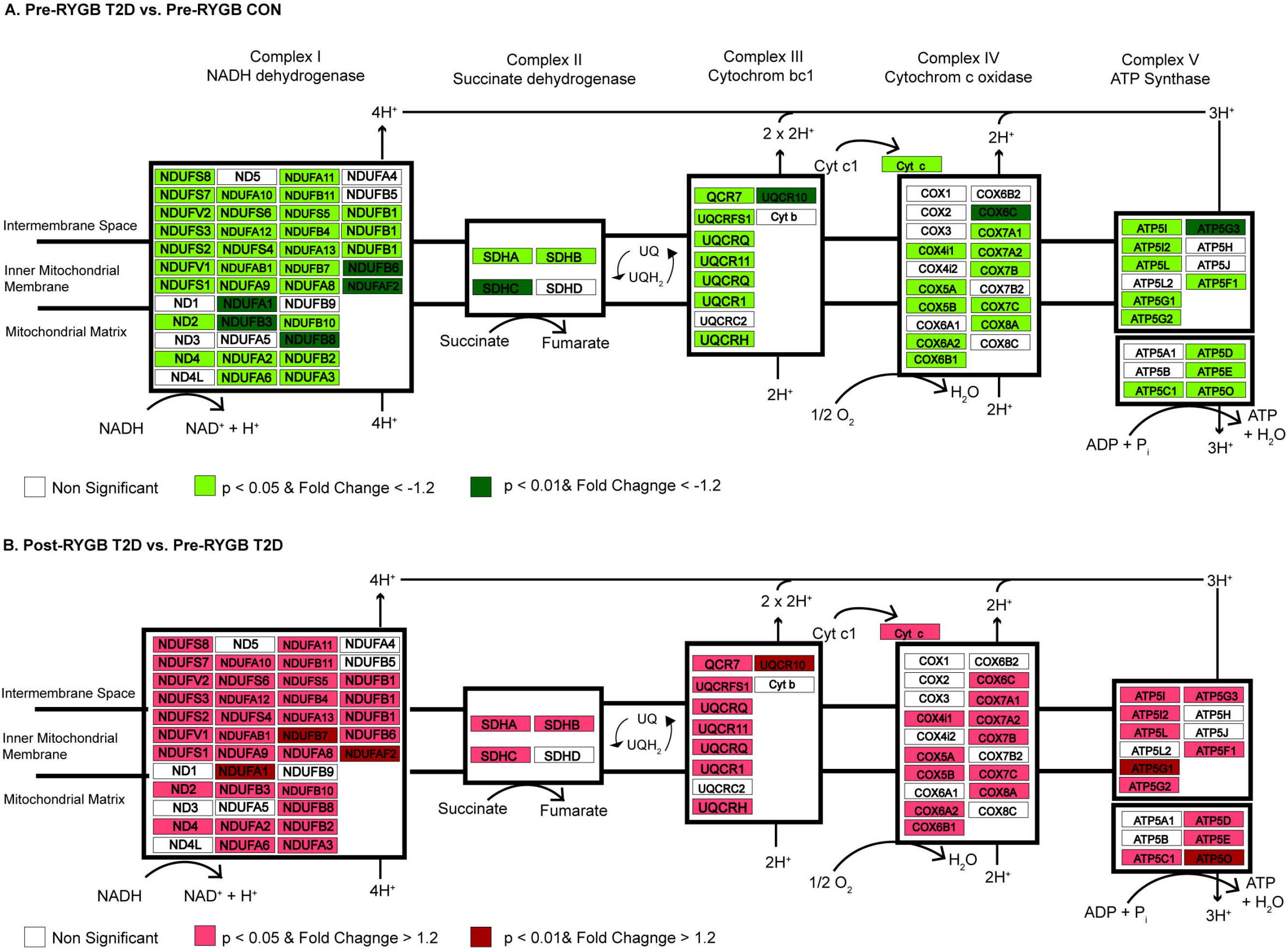
Differential regulation of genes of the mitochondrial complexes between individuals with and without diabetes and pre-to-post surgery in individuals with diabetes. **A)** Differentially regulated genes in Complex I, II, III, IV, and ATP Synthase in skeletal muscle of individuals with and without diabetes prior to surgery. Using an expanded gene list to include genes with p < 0.05, FC > −1.2 (lighter red shading; **Supplemental Table 6**), we show consistent down regulation of genes in mitochondrial complexes in individuals with diabetes prior to surgery. Green shading indicates p < 0.01 and FC > −1.2. **B.)** Differentially regulated genes in Complex I, II, III, IV, and ATP Synthase in skeletal muscle of individuals with diabetes pre-to-post surgery. Using an expanded gene list to include genes with p < 0.05, FC > −1.2 (lighter green shading; **Supplemental Table 6**), we show consistent up regulation of genes in mitochondrial complexes 1-year post surgery. Red shading indicates p < 0.01 and FC > 1.2.

We also show concurrent downregulation of key metabolic regulatory genes (**Table 2**) in individuals with diabetes prior to surgery. Hexokinase 2 (HK2; FC = −3.4), succinate dehydrogenase complex, subunit C (SDHC; FC = −3.1), isocitrate dehydrogenase 1 (IDH1; FC = −1.8), and pyruvate kinase muscle (PKM; FC = 1.5) were all downregulated in individuals with diabetes. Taken in conjunction with decreased gene expression of the mitochondrial complexes, Black women with T2DM show coordinated down regulation of aerobic metabolism genes in comparison to their non-diabetic counterparts.

### RYGB normalizes skeletal muscle gene expression profiles of Black women with diabetes

One year following RYGB the T2D group had significant improvement in clinical profiles similar to CON: normalized glycemia, reduced hyperinsulinemia, improved HOMA-IR, and significant reduction in excess weight and BMI. Comparison of skeletal muscle gene expression profiles one year following RYGB surgery between those with and without diabetes resulted in only 3 differentially regulated probes between the groups. This indicates that RYGB, and the following lifestyle changes not accounted for here, resulted in skeletal muscle gene expression of individuals with diabetes normalizing to their non-diabetic counterparts. This occurred despite the individuals with T2D being slightly older, suggesting that this age difference was likely not responsible for the initial differences in gene expression. Similarly, the normalization indicates the robust effects of the surgery/weight loss. Further comparison of skeletal muscle expression profiles pre-to-post RYGB in individuals with T2DM showed that almost all (43 of 48) differentially regulated genes increased following surgery and biological interpretation of these genes identified their role in mitochondrial function and aerobic metabolism.

We identified significant upregulation (Table 3) of mitochondrial complex I genes (NDUFB8, NDUFAF2, NDUFA1), ATP synthase subunits (ATP5O and ATP5G1), and cytochrome c oxidase subunit Vic (COX6C) in individuals with T2DM one-year post RYGB surgery. Skeletal muscle ETC protein content has been shown to be progressively diminished, including reductions in COX6C, in nondiabetic individuals with obesity and individuals with T2DM in comparison to lean counterparts (33). The degree to which these mitochondrial gene expression changes drive improved clinical phenotypes is indeterminable, but even changes in a small number of genes can have cascade like effects on cellular signaling and function. For instance, they may be heavily involved in promoting epigenetic (e.g. DNA methylation) changes in muscle (15; 34). Gene expression changes 52 weeks following surgery was more strongly linked to epigenetic (DNA methylation) changes in skeletal muscle than 2 weeks post-surgery when insulin resistance had already resolved (15). Barres et al. (34) also showed that changes in expression of three gene was linked to changes in over 100 CpG methylation sites. Thus, interpretation that gene expression changes in aerobic metabolism and mitochondrial function genes cannot be limited to improved mitochondrial function or flexibility driving improved phenotype (e.g. resolution of insulin resistance or T2DM). As with our pre-surgery analysis, expanded analysis (p < 0.05 and FC = |1.2|); Supplementary Table 6) shows consistent upregulation of a number of genes in all mitochondrial complexes one year following surgery in those with diabetes (**Figure 3B**).

### Expression of 3-hydroxybutyrate dehydrogenase, type 1 is reduced in skeletal muscle of Black women with T2D and improved following RYGB

Ketone bodies are important metabolic fuels produced in the liver and metabolized in the mitochondria of non-hepatic tissues (35). In our analysis we show reduced expression of 3-hydroxybutyrate dehydrogenase, type (BDH1), an important catalyst of ketone metabolism, in skeletal muscle of T2D (Table 2) and significantly upregulated one-year following RYGB (Table 3). Expression of BDH1 in adipose tissue has been positively correlated to insulin sensitivity in ~9,000 Finnish men (36), but not associated with circulating ketone levels. Expression of BDH1 in skeletal muscle is modulated by PGC-1α (35), but its potential as a significant modifier of skeletal muscle metabolic function following surgery has not been explored to date. Given the role β-hydroxybutyrate, the primary circulating ketone body, as a metabolic intermediate and its potential epigenetic signaling actions (37), more exploration in to this gene and protein in the context of weight loss surgery are warranted.

### Conclusion

In the present study, we explored skeletal muscle gene expression profiles of Black women with and without T2DM prior to and one year following RYGB. The presence of T2DM resulted in coordinated downregulation of genes in aerobic metabolism and oxidative metabolic pathway, specifically, subunits for mitochondrial complex I and complex II, as well as other key metabolic regulatory genes. One year following surgery, skeletal muscle gene expression profiles of Black women with T2DM had normalized to their non-diabetic counterparts, driven largely by improvements in genes involved in aerobic metabolism and mitochondrial function. Weight loss surgery is an incredibly effective treatment for weight loss and remission for T2DM, and changes in skeletal muscle gene expression may, in part, contribute to these phenotypic changes through improved metabolic and mitochondrial function as well as other cellular signaling functions such as epigenetic modifications. These genes and proteins should be further explored in the context of surgical weight loss and other lifestyle modification strategies to understand their role throughout the dynamic process of severe weight loss and improved metabolic phenotype.

## Acknowledgements

The authors would like to acknowledge the participants for their commitment to this research study.

## Supplemental Materials

Supplementary Table 1: Probes and statistics for the main effect of timepoint

Supplementary Table 2: Probes and statistics for the main effect of group

Supplementary Table 3: Probes and statistics for time x group interaction

Supplementary Table 4: Probes and statistics for baseline pairwise comparison (Pre-RYGB T2DM vs Pre-RYGB CON) of skeletal muscle gene expression

Supplementary Table 5: Probes and statistics for Pre-to-Post surgery skeletal muscle gene expression changes

Supplemental Table 6: Probes and statistics for expanded gene lists (p < 0.05) used biological pathway analysis

